# Schwann cell dysfunction contributes to diabetic wound pathology which is partially ameliorated by oncostatin M treatment

**DOI:** 10.64898/2026.02.24.707760

**Authors:** Shahreen Mahbub Rahman, Griffen Wakelin, Laura V. Young, JJ Parker, Leah Saleh, James Fawcett, Adam P.W. Johnston

**Author notes:** Current address for correspondence.

## Abstract

Chronic diabetic wounds represent a major clinical burden and are strongly associated with peripheral neuropathy, yet the contribution of nerve-associated Schwann cells to impaired healing remains poorly defined. Here, we investigated Schwann cell dynamics in cutaneous wound repair using the *db*/*db* model of type 2 diabetes. Full-thickness excisional wounds in *db*/*db* mice exhibited delayed closure, reduced dermal and epidermal thickness, and diminished cellular proliferation compared to non-diabetic controls. Diabetic wounds also demonstrated impaired re-innervation and a marked reduction in both total (S100β^+^) and dedifferentiated (p75NTR^+^) Schwann cells, including decreased Schwann cell proliferation. These findings indicate that diabetes disrupts the injury-induced Schwann cell response that is essential for normal repair. Transcriptomic analyses revealed that injury-activated Schwann cells upregulate multiple trophic factors, including oncostatin M (OSM), while single-cell RNA sequencing demonstrated broad expression of OSM receptors (*Osmr* and *Il6st*) across wound-resident keratinocytes, fibroblasts, and vascular-associated cells, suggesting widespread responsiveness to OSM signalling during repair. Therapeutic administration of OSM to diabetic wounds significantly accelerated closure, reduced wound width and area, and increased dermal and epidermal thickness. Mechanistically, OSM enhanced epidermal proliferation, angiogenesis, and cutaneous axon regeneration. Collectively, these data identify Schwann cell dysfunction as a contributor to impaired diabetic wound healing and demonstrate that augmenting a Schwann cell–derived paracrine signal can partially rescue key reparative processes. Our findings support a regulatory role for Schwann cells in coordinating epithelial, vascular, and neural repair responses and highlight OSM signalling as a potential therapeutic strategy for chronic diabetic wounds.

## INTRODUCTION

Skin wound repair is a dynamic and complex process generally consisting of the three overlapping phases, which include inflammation, proliferation and remodelling^1^. While normally efficient, wound healing is significantly impaired by conditions such as diabetes. These chronic non-healing skin wounds can develop into serious complications such as ulcers, which can ultimately lead to limb amputation^2^. Individuals with diabetes are very likely to be affected by a chronic wound at some point in their life, with estimates ranging from 8-35%^3,4^. While experimental therapies such as hyperoxygenation, stem cells, growth factors, and anti-inflammatory therapies are under investigation, none have proven universally effective across all patients^5^. The mechanisms underlying poor diabetic wound healing are not yet fully understood, though much research has focused on the roles of inadequate vascular supply^6^ and persistent inflammation^7^.

While not well understood, the presence of diabetic peripheral neuropathy (DPN), the painful dieback of nerve axons, especially in the hands and feet, is linked to a 7- to 20-fold higher risk of developing a foot ulcer in comparison with non-neuropathic diabetic individuals^8^. Nevertheless, the role of neuron/axon loss in diabetic wound healing has garnered relatively little attention despite substantial evidence supporting the essential involvement of nerves in normal wound repair^9,10^.

In this regard, while studies have highlighted the involvement of axon-derived neuropeptides such as substance P and calcitonin gene-related peptide (CGRP)^11^, our work has demonstrated that nerve-associated Schwann cells play a critical role in normal skin wound repair. We showed that injury to the skin also elicits significant local nerve damage and, as a consequence, activation and dedifferentiation of cutaneous Schwann cells into a proliferative and migratory phenotype^12,13^. These dedifferentiated Schwann cells dissociate from damaged nerves and migrate into the healing dermis, where they are located immediately adjacent to the dermal mesenchymal cells and facilitate their proliferation through paracrine actions^12,13^. Indeed, recent in vivo^14^ RNAseq^15^ and culture studies^13^ have shown that Schwann cells secrete ligands such as oncostatin M (OSM)^13^, PDGF-AA^13^, TGFβ^15^, and CCL2^14^ to promote the growth and self-renewal of dermal precursor cells^13^, formation of myofibroblasts^15^ and recruitment of monocyte-derived macrophages^14^ throughout various phases of wound healing. Importantly, ablation of Schwann cells or impairment of their actions through conditional knockout of regulatory genes such as Sox10^15^ or Sox2^12^ limits their activity and impairs the healing process. Taken together, these results highlight that Schwann cells are indispensable for normal wound repair; however, the impact of diabetes on Schwann cell activity remains poorly defined.

In the present investigation, we found that the quantity of total and dedifferentiated Schwann cells are dramatically reduced in the wounds of *db*/*db* diabetic mice, which demonstrate poor wound repair kinetics. Importantly, we revealed that treatment with the Schwann cell secreted factor OSM enhanced wound repair, cell proliferation, angiogenesis and innervation. Collectively, this work supports a regulatory role for Schwann cell dysfunction in chronic wounds and highlights the potential of Schwann cell secreted factors to influence multiple aspects of wound repair and axon regeneration.

## MATERIALS AND METHODS

### Animals and wounding

All experimental procedures were approved by the animal care committee of Dalhousie University and were performed according to the Canadian Council of Animal Care (CCAC) Guidelines for the Care and Use of Laboratory Animals. *db*/*db* and *db/+* control (B6.BKS(D)-Lepr^db^/J, strain 000697)^16^ were purchased from Jackson Labs at 8-12 weeks of age. For skin wound healing experiments, *db/db, db/+*, mice were anesthetized, the dorsal hair shaved, and the exposed skin cleaned with ethanol. Two 6-mm diameter full-thickness excisional wounds were administered on each side of the dorsal midline using a biopsy punch (Miltex) as described previously^12,17,18^. Mice were administered subcutaneous meloxicam for analgesia, individually housed, and wounds were left uncovered and tissues isolated at the time indicated. N = 5-7 mice per group per timepoint were utilized for experiments quantifying Schwann cells, innervation, EdU and wound healing characteristics between *db*/*db* and *db/+* mice. Wound morphometrics were tracked through photos at the times indicated. Wounds were excised, fixed in 4% PFA or 10% neutral buffered formalin and processed for histology, immunohistochemistry or immunofluorescence.

### Oncostatin M treatment experiments

For OSM treatment experiments, n=7 male 10-week-old *db*/*db* mice were administered bilateral 6-mm full-thickness wounds as described above. At the time of wounding, and on days 2, 4 and 7, one wound per animal received 200ng of OSM (R&D Systems, Minneapolis, MN) administered intradermally in PBS at three sites adjacent to the wound. The opposite wound received PBS administered at three sites around the wound. Wound morphometrics were tracked as above, and skin wounds were isolated, fixed 10% neutral buffered formalin and processed for histology, immunohistochemistry or immunofluorescence.

### Tissue preparation and immunostaining

For skin analyses, cryosections and paraffin sections were utilized as previously described^12,17^. Briefly, for cryosections, skin was fixed overnight in 4% PFA, cryopreserved in 30% sucrose for 1-3 days, then frozen in OCT (Tissue-tek) prior to sectioning at 18 µm and mounting on salinized glass slides. For paraffin sections, wounds were fixed in 10% neutral buffered formalin, embedded in paraffin, bisected in the caudocranial direction, and serial sections from the central portion of the wound were generated. For immunofluorescence staining, slides were blocked with 5% BSA with 0.3% Triton-X100 for 1 hour at room temperature, incubated with primary antibodies overnight, followed by secondary antibodies for 2 hours at room temperate then counterstained with DAPI (Sigma). For paraffin sections, antigen retrieval was performed by submerging sections in tri-sodium citrate buffer (pH 6.0) for 20 min at 95°C. When paraffin sections were stained using immunofluorescence, endogenous autofluorescence was blocked utilizing the Tru-View autofluorescence blocking reagent (Vector) as per the manufacturer’s instructions. For colorimetric staining of paraffin sections, primary antibodies were detected using the ABC kit and visualized using a DAB kit (Vector Laboratories). Sections were visualized using a Zeiss AxioObserver 7 inverted microscope (Carl Zeiss Microscopy, Jena, Germany). The following antibodies were utilized for immunostaining: rabbit anti-p75NTR (1:500; Promega), rabbit anti-S100β (1:500; Dako), Rabbit anti-Ki67 (Abcam ab16667), rabbit anti-ßIII-tubulin (Abcam ab18207), goal anti-CD31 (R&D Systems AF3628). Secondary antibodies were anti-rabbit, or goat, Alexa-488 or 555 (1:1000; Molecular Probes).

### Quantification of wound cell types

To quantify Schwann cells within healing skin, tissue sections were immunostained for S100β or p75NTR, counterstained with DAPI, and manually counted in a blinded manner using Fiji (ImageJ), with values normalized to dermal wound area. βIII-tubulin–positive axons, CD31-positive capillaries, and Ki67-positive dermal and epidermal cells were quantified using the same approach as we have done before^12,17,18^. For each marker, two sections per sample were quantified, then averaged prior to statistical analysis. To identify proliferating Schwann cells, mice were intraperitoneally injected with 100mg/kg 5-ethynyl-2’-deoxyuridine (EdU) (Molecular Probes) 1 day before sacrifice, cryosections were generated and immunostaining for S100β. EdU was detected using the Click-iT Alexa Fluor 488 or 555 imaging kit (Thermo Fisher) as per the manufacturer’s instructions. The quantity of S100β^+^/EdU positive cells within the entire wound bed was manually calculated using ImageJ analysis software. The experimenter was blinded to the groups during analysis. Imaging was performed using a Zeiss Axio Observer 7 inverted microscope.

### Morphometric analyses

To quantify wound size, topographical digital photographs (Canon) were taken at the times indicated post-wounding. A ruler was used to calibrate the images, and the wound margins were manually outlined using Fiji (ImageJ). Morphometric analyses of wound parameters were performed on hematoxylin and eosin-stained paraffin tissue sections from the central portion of the wound bed as described previously^12,17,18^. Wound width was measured as the distance between the wound margins, as defined by the last hair follicles. Wound area encompassed newly healed dermal and epidermal tissue, and epidermal and dermal thickness were measured at three points in the newly healed dermis/epidermis, then averaged. Imaging was performed using a Zeiss Axio Observer 7 inverted microscope, and analyses were performed in a blinded manner.

### Single-cell and bulk RNA sequencing analysis

For bulk RNA-seq analysis, raw data (ENA, PRJEB22372)^15^ were aligned to the GRCm38 reference assembly and quantified using subread^19^. Differential expression tests were performed in R (4.5.0) using the DESeq2 package (1.48.2)^20^. For visualization, only variance-stabilizing transformed values are shown. For single-cell RNA-seq analysis (scRNAseq), data (GSE142471)^21^ were processed using scanpy (1.12)^22^. Cells were normalized, principal component analysis was performed, and tSNE embeddings were used for visualization. The Wilcoxon rank-sum test was used for all statistical analyses. Cells with a total number of counts or genes greater than 3 median absolute deviations from the mean, or greater than 20 percent of counts mapping to mitochondrial genes, were excluded from analysis.

## STATISTICAL ANALYSIS

A repeated measures, 2-way analysis of variance (ANOVA) was used to analyze differences in topographical wound closure rate between *db*/*db* and *db/+* mice. A 2-way ANOVA was utilized to assess for differences in the number of S100β+ or p75NTR+ Schwann cells and extent of reinnervation (BIII tubulin+) following wounding between *db*/*db* and *db/+* mice. A repeated-measures 2-way ANOVA was used to analyze differences in topographical wound closure rates between OSM- and PBS-treated wounds. A two-tailed, paired Student’s t-test was utilized to examine the impact of OSM treatment on wound healing parameters (wound width, area, dermal thickness, epidermal thickness), number of Ki67+, CD31+ or BIII-tubulin+ cells. Significance was set at p < 0.05, and data are represented as mean ± standard deviation.

## RESULTS

### db/db diabetic mice demonstrate characteristic impairments in wound healing kinetics

To understand the extent of impaired wound healing with diabetes, we administered 6-mm full-thickness punch wounds to the dorsal skin of control (*db/+*) and *db*/*db* diabetic mice. *db*/*db* leptin receptor-deficient mice represent a model of type 2 diabetes, which develop obesity, hyperglycemia, hyperinsulinemia, and many diabetic complications, including DPN and impaired healing kinetics^23,24^. Tracking wound morphometrics throughout the healing time course demonstrated significant impairments in wound healing rate, evident by 5 days post-injury (Figure 1A, B). Control skin was largely healed by 10 days post-wounding, while some *db*/*db* wounds were still evident up to 21 days post injury (Figure 1B), in agreement with previous reports^23^. This was also evident in the analysis of H&E-stained (Figure 1C) cross-sections of 10-day wounded skin, which revealed significantly (p < 0.05) larger wound width and wound areas in *db*/*db* mice (Figure 1D, E) while dermal and epidermal thickness (Figure 1F, G) were reduced in comparison to control animals. Analysis of EdU incorporation (Figure 1H), as a measure of cell proliferation within the wound bed, demonstrated that diabetes significantly (p < 0.05) impaired dermal cell proliferation in *db*/*db* wounds in comparison to control wound samples (Figure 1I). Taken together, these results demonstrate that diabetes significantly impairs wound healing rate and reduces cell proliferation in healing tissue.

**Figure 1.**
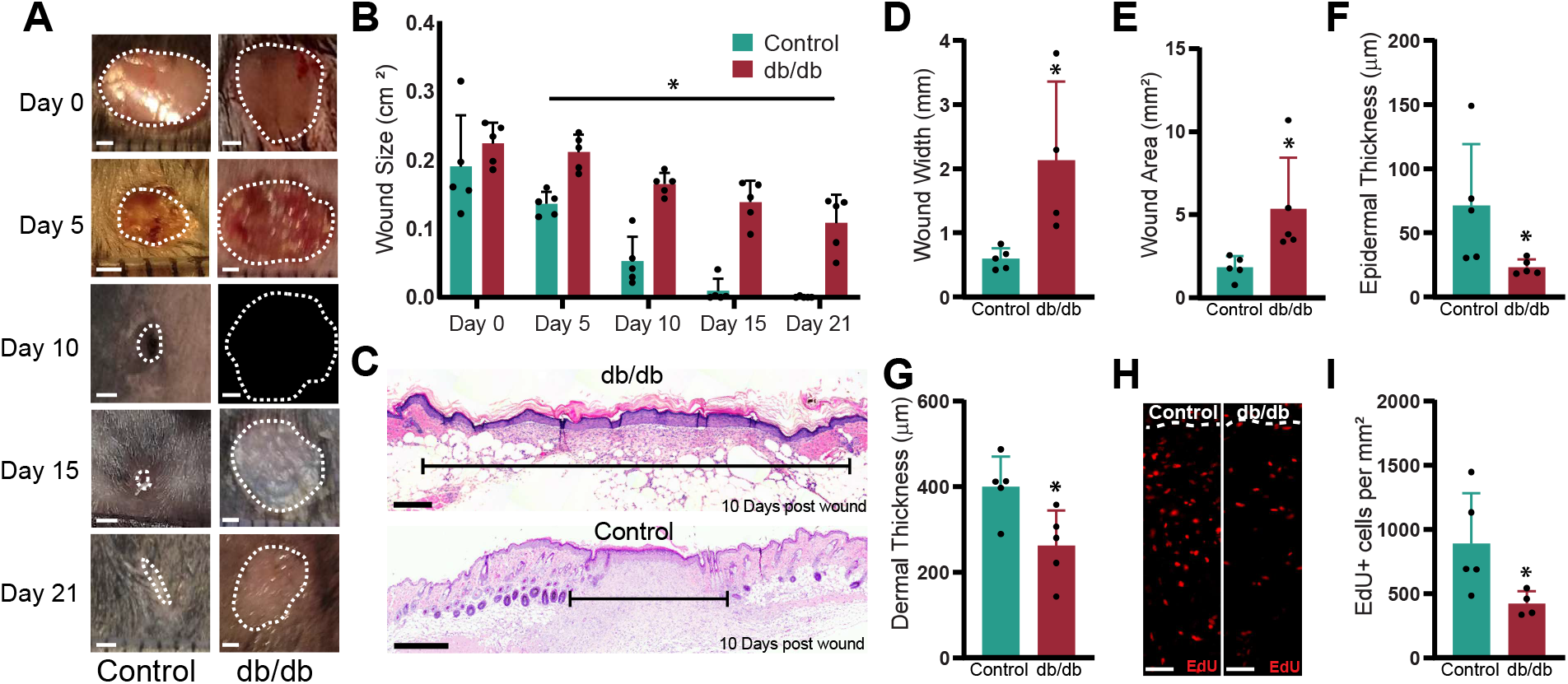
Diabetes impairs cutaneous wound healing. *db*/*db* and control *db*/*+* mice were wounded with a 6-mm punch biopsy, and wounds were photographed (A), and size was analyzed across time (B), demonstrating significant impairments in wound healing kinetics. Analysis of H&E staining (C) in 10-day post-injury skin for wound width (D), wound area (E), epidermal thickness (F) and dermal thickness (G) in *db*/*db* and *db/+* control mice, demonstrating significant decrements in wound healing capacity in *db*/*db* mice. Fluorescent images (H) and analysis (I) of cell proliferation as indicated by EdU in *db*/*db* and *db/+* control mice 10 days post-wounding, showing impaired dermal cell proliferation in *db*/*db* mice. * Indicates a significant p < 0.05 difference compared to control *db/+* mice. Scale bars: A = 1 mm, C = 500 µm, H = 50 µm.

### Diabetic wounds are neuropathic and lack Schwann cells

Next, to understand how impaired wound healing with diabetes relates to Schwann cell activity, we administered 6-mm full-thickness punch wounds to control (*db/+*) and *db*/*db* mice. First, we quantified the extent of wound re-innervation through βIII-tubulin immunostaining, which revealed a significant (p < 0.05) reduction in the density of dermal nerve axons at 10-days post wounding (Figure 2A-C). Further, immunostaining for p75NTR, a marker of dedifferentiated Schwann cells, revealed many “activated” nerve bundles containing EdU-positive proliferating, dedifferentiated Schwann cells in proximity to healing wounds 10 days post-injury in control wounds, which were largely absent in *db*/*db* diabetic wounds (Figure 2J-M). Next, we examined the quantity of Schwann cells within and around the wound beds of control and diabetic mice at 10 and 15 days post-wounding using immunostaining. This demonstrated that in control mice, many S100β-positive (143.3 ± 56.2 mm^2^) and p75NTR-positive (237.4 ± 68.7 mm^2^) Schwann cells were found intermixed with dermal cells within the healing skin (Figure 2E, K), in agreement with our previous reports^12,13^. However, by comparison, *db*/*db* mice demonstrated significant (p < 0.05) reductions in the number of S100β-positive (15.6 ± 10.4 mm^2^) and p75NTR-positive Schwann cells (37.1 ± 14.6 mm^2^) post-wounding, which was also evident at 15 days post-wound (Figure 2D, H, J, N). Diabetes appeared to negatively impact Schwann cell activity as there was a significant reduction (p < 0.05) in the quantity of S100β-positive cells, co-expressing EdU in the wound beds of diabetic mice (Figure 2F, G, I). Collectively, these results demonstrate that Schwann cell content and activity are diminished in diabetic wounds, coincident with poor wound healing and innervation.

**Figure 2.**
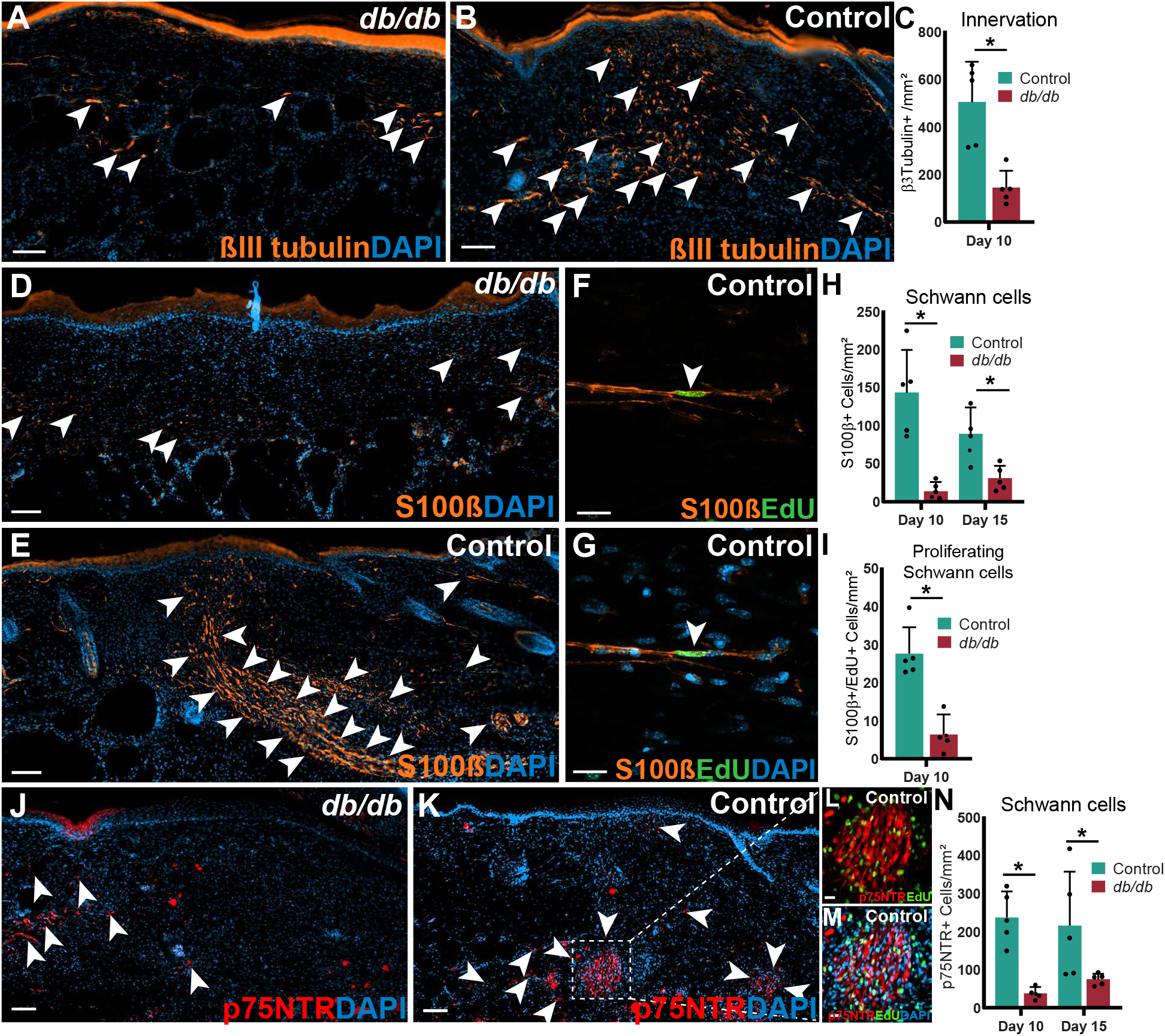
Diabetes reduces Schwann cell quantity and axon density in wounded skin. *db/db* and control *db*/+ mice were wounded with a 6-mm punch biopsy, and wounds were isolated at 10 and 15 days post-injury. Immunofluorescent staining of *db/db* (A) and control (B) wounds with quantification (C) of βIII-tubulin-positive axons (orange) demonstrating a significant impairment in wound innervation in *db/db* mice. Immunofluorescent staining of S100β (orange, D-G) and EdU (green, F, G) in *db/db* (D) and control (E-G) skin demonstrates significant reductions in the total number (H) and proliferating (I, EdU-positive), Schwann cell populations in *db/db* wounds. Immunofluorescent staining of *db/db* (J) and control (K) wounds with quantification (N) of p75NTR-positive cells (red) demonstrating a significant reduction in dedifferentiated Schwann cells in *db/db* mice. High-magnification images of p75NTR (red) and EdU (green, L, M) from panel K demonstrating the presence of activated nerve bundles in control skin. * Indicates a significant p < 0.05 difference compared to control *db*/+ mice. Arrowheads indicate positive cells. Scale bars: A, B, D, E, J, K = 100 μm; F, G, L, M = 20 μm

### OSM signalling is upregulated during skin wound repair

Schwann cells are critical to the healing of healthy skin^12,17^, through the paracrine secretion of trophic factors. Previous work has demonstrated that dedifferentiated Schwann cells display elevated expression of many growth factors and cytokines. As such, a reduction in Schwann cell quantity may impair diabetic wound healing due to insufficient trophic support. To help identify secreted factors that are upregulated in Schwann cells during wound healing *in vivo*, we analyzed published RNAseq data^15^ of Schwann cells, FACS purified from wild-type (WT), uninjured and wounded skin. This revealed significant (p < 0.05) increases in the expression of many growth factors with known roles in tissue repair and remodeling such as platelet-derived growth factor c (*Pdgfc*), leukemia inhibitory factor (*Lif*), *Ccl2* and *Osm* (Figure 3A-D). We focused on OSM, a member of the interleukin-6 (IL-6) cytokine family, as our prior studies demonstrated that OSM enhances dermal precursor cell proliferation and, when ectopically delivered to regenerating digit tips, promotes bone growth and rescues denervation-induced defects in digit regeneration^17^. Next, to assess the potential relevance of OSM in wound repair, we examined published scRNAseq data of uninjured and wounded skin^21^ (Figure 3E-F). Clustering analysis revealed characteristic skin cell types in each condition (Figure 3E), including keratinocytes, fibroblasts, immune cells and vascular-related cell types such as smooth muscle cells and pericytes (Figure 3F). Next, to identify which cell types OSM may act upon, we examined the expression of the OSM receptors *Osmr* and *Il6st*, which were widely expressed in populations, including keratinocytes, fibroblasts and vascular cells under basal conditions and following injury (Figure 3G-J). Additionally, wounding elicited significant upregulations (p < 0.05) of *Osmr* and *il6st* in subsets of keratinocytes, pericytes/smooth muscle cells and fibroblasts (Figure 3I-J).

**Figure 3.**
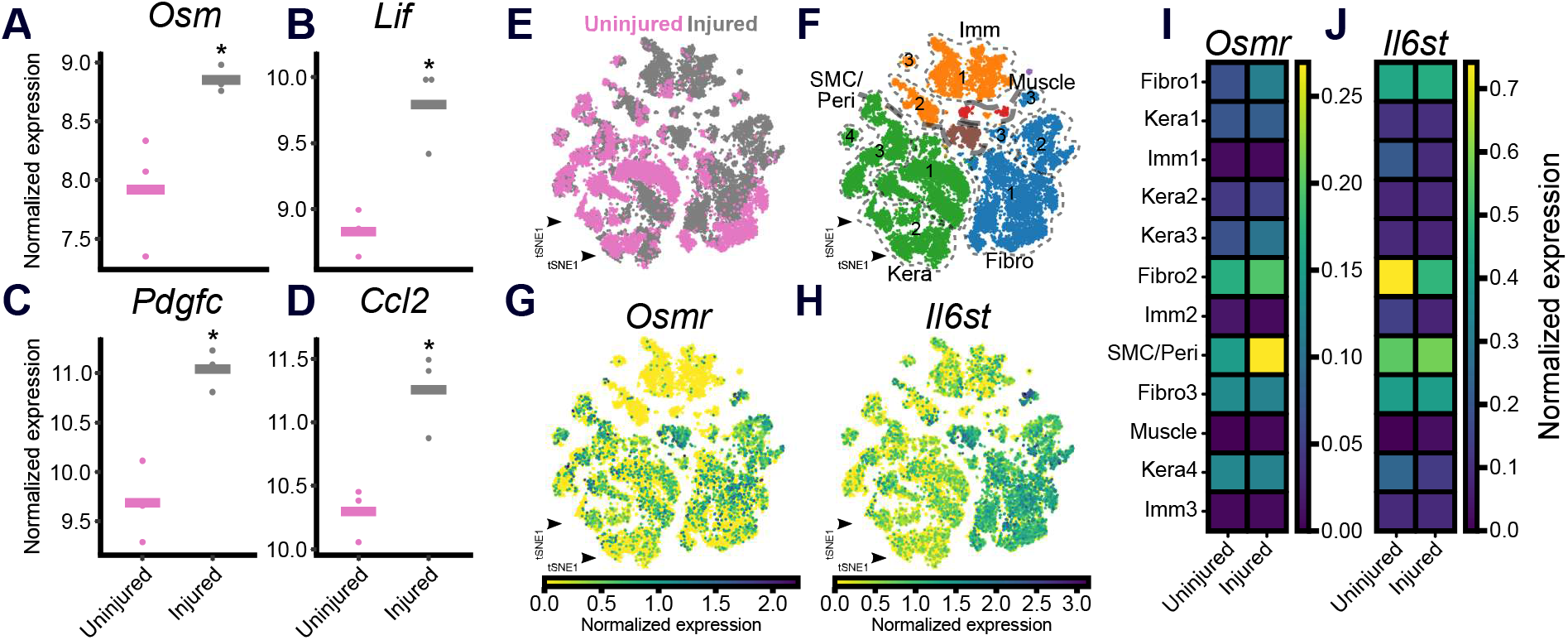
OSM signalling is active during wound repair. Bulk RNA sequencing analysis of Schwann cells FACS purified from uninjured skin and wounds administered 7 days earlier, demonstrating significant upregulations in the expression of *Osm* (A), leukemia inhibitory factor (*Lif*, B), platelet-derived growth factor C (*Pdgfc*, C), and *Ccl2* (D) in Schwann cells following injury. UMAP plots of single-cell RNA sequencing (scRNAseq) analysis demonstrating cell clusters in injured and uninjured skin (E), including clusters of keratinocytes, fibroblasts, immune cells and vascular-related smooth muscle cells and pericytes (F). Feature plots (G, H) and matrix plots (I, J) demonstrating the expression of the OSM receptors *Osmr* and *Il6st* in wound resident cell types and their increased expression in select clusters of fibroblasts, vascular cells and keratinocytes (I, J). Analysis in A-D is from Parfejevs et al.^15^ while E-J is from Haensel et al.^21^, respectively.

### OSM treatment accelerates diabetic wound healing

These findings identify OSM as a candidate molecule capable of signalling multiple wound-resident cell types, with the potential to modulate the wound healing response. To test this, we performed bilateral 6-mm punch wounds to *db*/*db* mice and applied either OSM or vehicle (PBS) to each wound, intradermally, on days 0, 2, 4 and 7 and wounds were isolated on day 9 and examined histologically (Figure 4A). Quantification of the mean wound size demonstrated that OSM treatment had a clear impact on skin repair, and by 4 days post-wound, OSM-treated wounds were on average 18% smaller and by 9 days were 45% smaller in area (Figure 4B, C). These findings are consistent with analysis of H&E-stained tissue sections, which showed that OSM treatment significantly (p < 0.05) reduced mean wound width by 33% (OSM, 2.89 ± 0.80 mm vs. Con, 4.36 ± 1.0 mm) and wound area by 43% (Con, 2.10 ± 0.85 vs. OSM, 1.18 ± 0.42 mm^2^, Figure 4E, F), while significantly (p < 0.05) increasing both dermal and epidermal thickness (Figure 4G, H).

**Figure 4.**
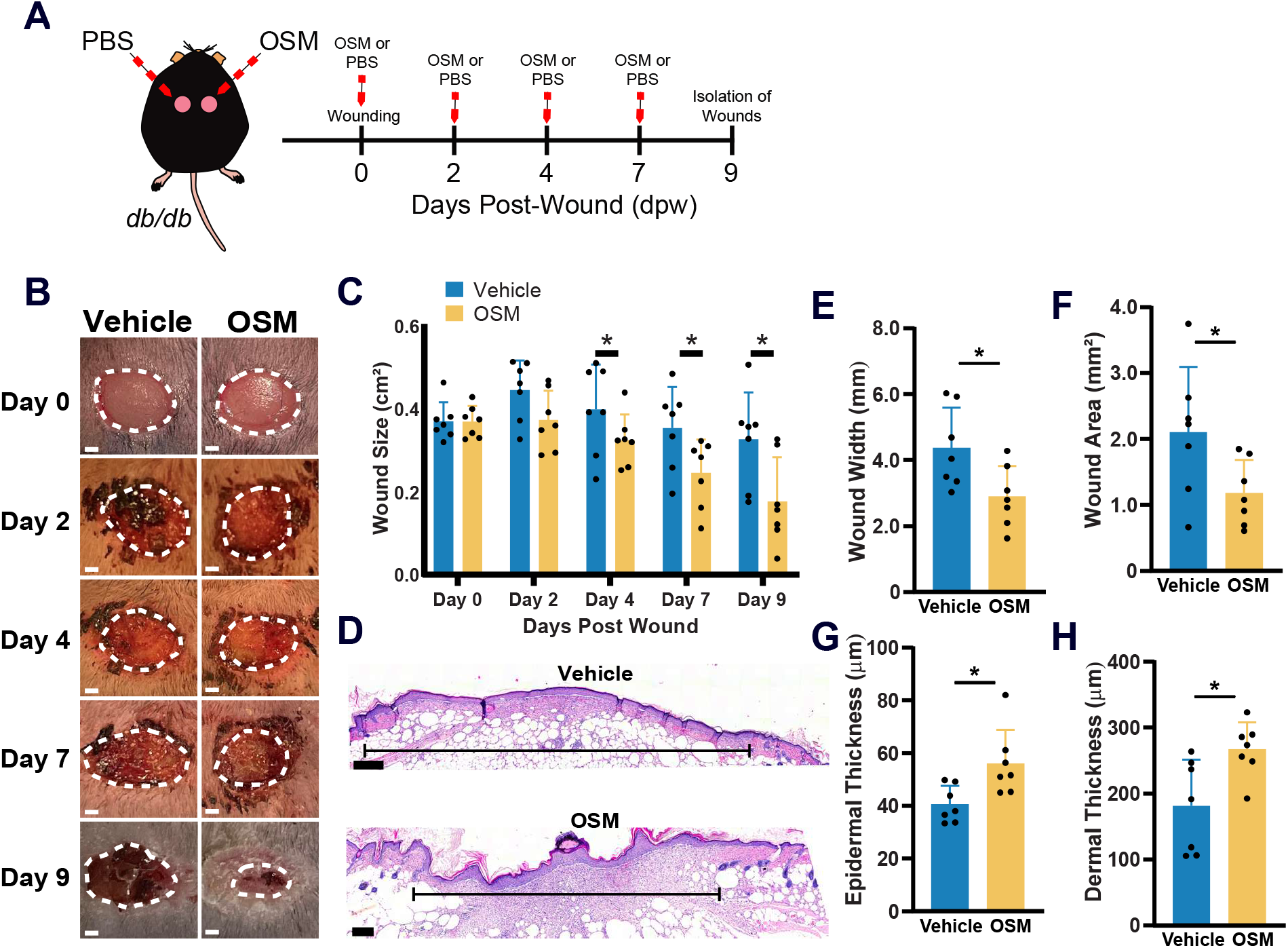
OSM treatment improves wound healing in *db*/*db* mice. (A) Schematic of the experimental design where *db*/*db* mice were wounded and administered vehicle or 200ng of OSM on days 0, 2, 4, 7 post-wound. Images (B) and analysis (C) of vehicle and OSM-treated wounds demonstrating a significant acceleration in wound healing kinetics with OSM treatment. Analysis of H&E staining (D) of vehicle and OSM-treated wounds for wound width (E), wound area (F), epidermal thickness (G) and dermal thickness (H), demonstrating enhanced wound healing parameters. * Indicates a significant p < 0.05 difference compared to vehicle treatment. Scale bars: B, D = 500 µm.

### OSM enhances cell proliferation, angiogenesis and axon regrowth

Next, we examined the mechanisms by which OSM enhanced wound healing. Immunostaining for the proliferation marker Ki67 demonstrated that, in comparison to vehicle-treated wounds, OSM treatment significantly (p < 0.05) increased the number of proliferating epidermal cells (Con, 21.7 ± 17.6 vs. OSM, 55.8 ± 23.4 cells/mm^2^) while dermal proliferation was similar between groups (Figure 5A-D). Additionally, we examined wound bed angiogenesis through immunostaining for the endothelial cell marker CD31. This revealed a significant (p < 0.05) increase in the quantity of CD31 cells within OSM-treated wounds in comparison to controls (Con, 100.7 ± 75.5 vs. OSM, 164.7 ± 102.0 vessels/mm^2^, Figure 5E-G). Finally, we examined wound bed innervation, which demonstrated that OSM significantly (p < 0.05) increased the density of βIII-tubulin-positive dermal axons when compared to vehicle treatment (Con, 0.0019 ± 0.0012 vs. OSM, 0.0037 ± 0.0015 βIII-tubulin area (µm^2^)/total wound area (µm^2^), Figure 5H-J). Taken together, these results show that administration of OSM to the wounds of *db*/*db* diabetic mice accelerates wound healing, cell proliferation, angiogenesis and axon regeneration.

**Figure 5.**
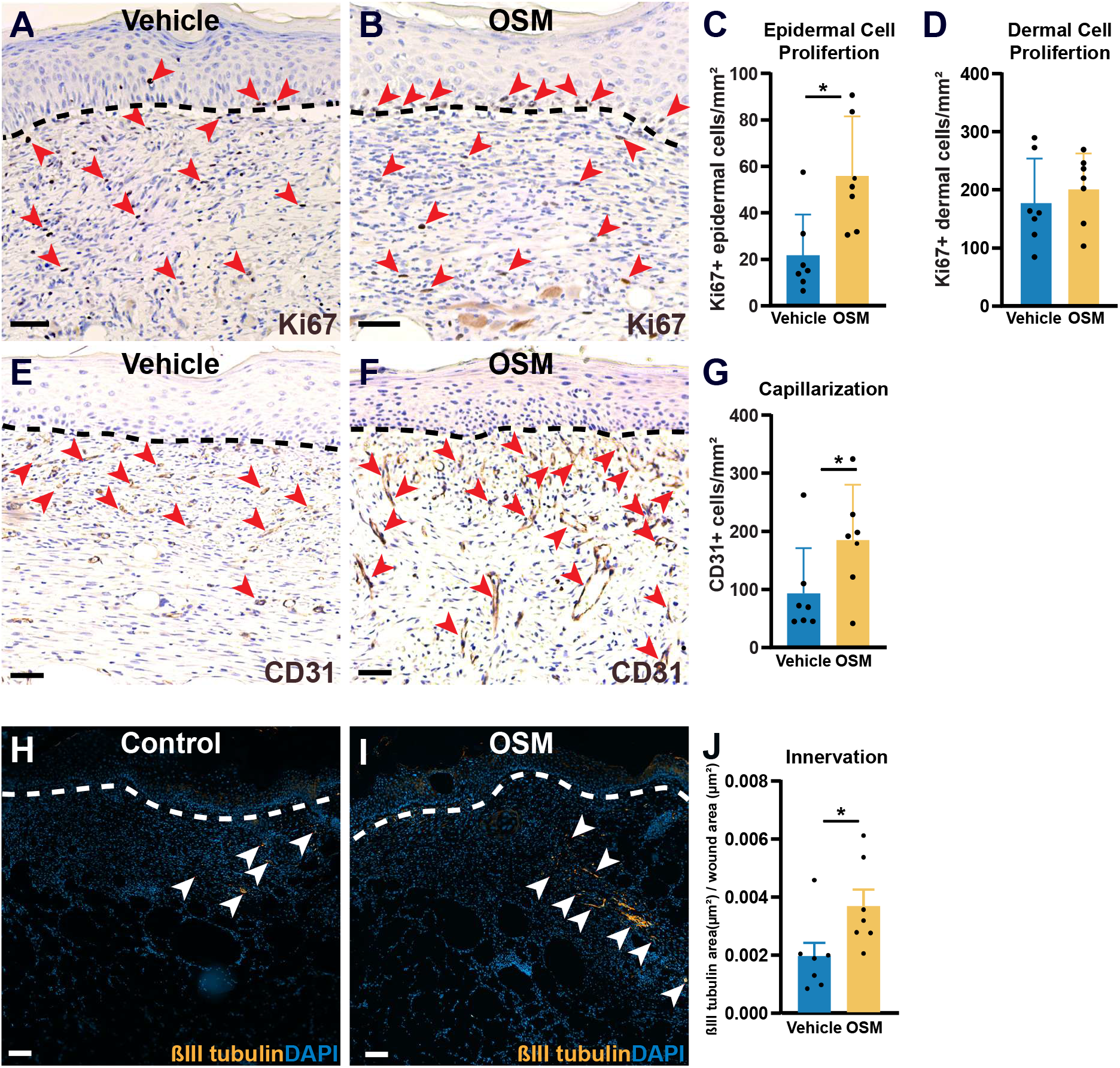
OSM enhances wound site angiogenesis, innervation and cell proliferation. Immunohistochemical staining (A, B) and quantification of Ki67-positive proliferating epidermal (C) and dermal (D) cells in vehicle and OSM-treated wounds of *db*/*db* mice, demonstrating greater epidermal cell proliferation with OSM treatment. Immunohistochemical staining (E, F) and quantification (G) of CD31-positive capillary endothelial cells in wounds of *db*/*db* mice demonstrating a significant increase in wound angiogenesis with OSM treatment. Immunofluorescent staining (H, I) and quantification (J) of βIII-tubulin-positive nerve axons showing greater wound bed innervation with OSM treatment in *db*/*db* mice. Scale bars: A B, E, F = 50 μm, H, I = 200 μm. * Indicates a significant p < 0.05 difference compared to vehicle treatment.

## DISCUSSION

In the current investigation, we demonstrate that 1) Schwann cells, which play an integral role in the healing of normal skin wounds, are reduced and impaired in the wounds of *db*/*db* diabetic mice, 2) OSM receptors are widely expressed on several wound-resident cell types including keratinocytes and vascular cells and 3) treatment with OSM, which is known Schwann cell factor, accelerates the healing of diabetic wounds and correspondently enhances epithelialization, angiogenesis and axon regeneration. Taken together, these results reinforce the role of Schwann cells in skin wound repair and suggest that OSM is a candidate molecule for clinical translation to enhance the closure of chronic non-healing diabetic skin wounds.

The current results support the engagement of Schwann cells in the process of normal wound repair, whereby skin injury engages a characteristic activation and dedifferentiation of Schwann cells as they migrate from their nerve-resident environment into the healing dermis. This is in agreement with our previous work^12,17^, the research of Dr. Sommer’s group^14,15^ and others^25,26^. It is now appreciated that this Schwann cell response is a prerequisite of normal wound repair, and genetic perturbation of Schwann cell activity results in aberrant wound healing^12,14,15,17^. In this regard, we describe that *db*/*db* diabetic wounds, which show impaired healing characteristics, contain fewer and less proliferative Schwann cells. These results are in line with studies that observed reduced dedifferentiated Schwann cell content in the wounds of *db*/*db*^26^ mice and streptozotocin-treated mice, a model of type 1 diabetes^25^. While it is clear that the diabetic milieu is detrimental to Schwann cell activity, the upstream signals that drive cutaneous Schwann cell dysfunction are poorly defined. Indeed, reactive oxygen species, advanced glycation end products and alterations in metabolism have all been implicated in Schwann cell dysfunction with diabetes^27^. Defining the specific pathways involved and identifying additional regulatory mechanisms will be essential for developing targeted therapies to restore Schwann cell function.

Based on their known paracrine actions, these results highly suggest that reductions in Schwann cell content are likely to impair the necessary trophic support for dermal and epidermal proliferation during wound repair. However, direct evidence demonstrating a causal role for Schwann cells in diabetic wound healing, rather than a correlative association, remains lacking. Experimental approaches such as Schwann cell transplantation or targeted molecular correction of intrinsic Schwann cell defects would more conclusively establish their functional contribution to diabetic wound pathology. Furthermore, additional studies are needed to establish whether a similar relationship exists in human clinical wound samples. In this context, there is a pressing need to define the role of Schwann cells in other cutaneous pathologies, including venous and arterial ulcers as well as burn injuries, to fully delineate the scope of Schwann cell function in clinical wound biology.

We previously defined that Schwann cells secrete OSM and that treatment of regenerating digit tips with OSM promotes bone regrowth and rescues denervation-induced defects in digit tip regeneration^17^. Based on this, we identified potential cellular targets of OSM signalling during wound repair, which revealed that the OSM receptors are widely expressed and dynamically regulated during the repair process. Furthermore, we demonstrate that OSM accelerates wound healing in diabetic mice by enhancing multiple reparative processes, including epidermal proliferation, angiogenesis, and axonal regrowth. These findings support those of Das and colleagues^28^ who demonstrate that active OSM is deficient in diabetic skin wounds and that exogenous treatment with OSM enhances re-epithelization. Additionally, Ishida et al^29^ reported that the expression of OSM and OSMR are reduced in the wounds of diabetic mice and that OSMRβ null mice demonstrate wound healing impairments, which accompany angiogenic defects.

Less is known about the role of OSM in peripheral neuron biology, and to our knowledge, this is the first report demonstrating that OSM can promote cutaneous axon regrowth during wound healing of diabetic mice. Recent work suggests that OSM signalling is involved in itch sensation, and cutaneous administration of OSM to uninjured mice elicits sensory neuron lengthening and hypersensitivity to mechanical stimuli^30^. Future work should examine the potential of OSM to enhance axon regeneration following neuronal injury or to preserve axon integrity under conditions of DPN.

Taken together, our work supports the expanding role of dedifferentiated Schwann cells in efficient wound repair and identifies their diminished response as a potential mediator of diabetic wound healing deficits. OSM, a multifunctional cytokine known to be secreted by Schwann cells, enhances wound healing in diabetic mice and has the potential to enhance angiogenesis, epithelial proliferation and axon regeneration.

## ACKNOWLEDGEMENTS

This work was supported by a CIHR Project Grant (141926), JF is supported by an NSERC Discovery Grant, SMR was supported by the Canada Graduate Scholarship (CGS-M), and LVY was supported by the Jeanne and J.-Louis Lévesque scholarship.

## AUTHOR CONTRIBUTIONS

SMR - Conceptualization, Investigation, Methodology, Analysis, Writing; GW - Investigation, Methodology, Analysis, Writing. LVY - Conceptualization, Investigation, Methodology. JJP - Conceptualization, Investigation, Methodology, Analysis. LS - Methodology, Analysis. JF - Conceptualization, Investigation, Methodology, Writing, Editing. APWJ Conceptualization, Investigation, Methodology, Analysis, Writing, Editing, Supervision, Funding acquisition.

## Notes

### Competing Interest Statement

The authors have declared no competing interest.

### Summary of Updates

The author affiliations have been updated.

